# Androgen receptor binding sites are highly mutated in prostate cancer

**DOI:** 10.1101/225433

**Authors:** Tunç Morova, Mehmet Gönen, Attila Gursoy, Özlem Keskin, Nathan A. Lack

**Affiliations:** School of Medicine, Koç University, Istanbul, Turkey; College of Engineering, Koç University, Istanbul, Turkey; Vancouver Prostate Centre, University of British Columbia, Vancouver, Canada

**Keywords:** Whole Genome Sequencing, Prostate Cancer, Androgen Receptor, Transcription Factor, Mutational Signature

## Abstract

Androgen receptor (AR) signalling is essential to nearly all prostate cancer cells. Any alterations to AR-mediated transcription can have a profound effect on prostate carcinogenesis and tumour growth. While the AR protein has been extensively studied, little is know about mutations to the non-coding regions where AR binds to DNA. Using clinical whole genome sequencing, we demonstrate that AR binding sites have a dramatically increased rate of mutations that is greater than any other transcription factor and specific to only prostate cancer. Demonstrating this may be common to lineage-specific transcription factors, estrogen receptor binding sites had an elevated rate of mutations in breast cancer. Based on the mutations observed at the binding site of AR and other related transcription factors, we proposed that AR occupancy impairs access of base excision repair enzymes to endogenous DNA damage. Overall, this work demonstrates that non-coding AR binding sites are frequently mutated in prostate cancer and may potentially act as driver mutations.

## INTRODUCTION

Cancer arises through the sequential accumulation of mutations that induce neoplastic transformation and uncontrolled proliferation. Each mutation can provide remarkable insight into the history of the cancer with different types of mutations arising from different DNA damage [1]. Somatic mutations do not occur in a normal distribution across the genome and are affected by several variables including GC content, replication time, distance to telomere and chromatin compaction [2–4]. Recent studies have demonstrated that transcription factor (TF) binding to DNA can also correlate with a higher rate of mutations [5,6]. Elegant work combining XR-seq from UV-treated skin fibroblast cells and large-scale whole genome sequencing (WGS) demonstrated that in skin cancer, TF binding impairs nucleotide excision repair machinery (NER) [7]. By physically preventing the access of repair enzymes, TF binding causes a higher rate of UV-mediated mutations in skin cancer. However, it is unlikely that only NER is affected by TF binding given the diversity of mutations observed in different cancer types.

Prostate cancer (PCa) is an extremely common disease that affects an estimated one out of every seven North American men in their lifetime. At all stages of PCa development, androgen receptor (AR) mediated transcription is critical to the growth of the tumour. Following activation, the AR translocates from the cytoplasm to the nucleus where it interacts with pioneer factors such as FOXA1 or HOXB13 before binding to chromatin. The vast majority of AR binding sites (ARBS) are located in intronic or intergenic regions [8,9]. Many of the ARBS contain an androgen response response elements (AREs) that consists of a DNA 15-bp palindromic sequence containing two hexameric 5’-AGAACA-3’ half sites arranged as an inverted repeat with a 3bp spacer [10,11]. Once bound to DNA, the AR recruit’s various coactivators that eventually initiate transcription of pro-mitotic genes. Several factors have been demonstrated to affect AR-mediated transcription such as epigenetic modifications, pioneer factors and chromatin accessibility [9,12]. Demonstrating the importance of these co-activtors and pioneer factors, HOXB13, GATA2 and KDM1A have been shown to be critical for AR signalling and are required for the growth of PCa cell lines [13–16].

In addition to initiating PCa growth, there is also evidence that AR signalling is associated with DNA damage. Goodwin *et al*. demonstrated a feedback loop where DNA repair genes activate the AR upon DNA damage and subsequently promotes DNA repair [17]. Further, AR has itself been demonstrated to induce double stranded breaks (DSB) via topoisomerase IIb (TOP2B) [18]. Specifically, AR recruits TOP2B to introduce DSB that relax torsional stress and allow transcription. These DSB are not typically recombinogenic and can be repaired by DNA repair mechanism. However, additional genotoxic stress can prevent repair of these DSBs and increase the rate of breaks by activation induced cytidine deaminases or LINE-1 repeat-encoded ORF2 endonucleases, thereby leading to structural variations such as the common TMPRSS2:ERG fusion [19].

There has been extensive research to identify protein coding “driver” mutations in both primary and castrate resistant PCa [20,21]. From these large studies numerous deletions (PTEN, CADM2), structural variants (ETS fusions) and single nucleotide variations (FOXA1, SPOP) have been identified as likely driver mutations in primary PCa. However, until recently the impact of non-coding mutations has been poorly understood. This is changing as their importance is increasingly becoming more evident. One of the first non-coding driver mutations identified, was found at the promoter of telomerase reverse transcriptase (TERT) [22,23]. Due to this non-coding mutation, expression of *TERT* is upregulated and causes an increased repair of shortened telomeres [24]. In PCa, a non-coding mutation to a polymorphic regulatory element was identified that impacted the regulation of DNA repair and hormone-regulated transcript levels in SPOP mutant patients [25]. Given their potential role in modifying the regulatory landscape of PCa, a better understanding of non-coding mutations is critical for more effective treatments.

While AR has been previously shown to induce DNA damage *in vitro*, the relatively low frequency of somatic mutations in PCa (~1 SNV/Mb) has prevented the study of TF-mediated DNA damage in clinical samples. Therefore, using large-scale WGS data we investigated how TF binding affects somatic mutations in PCa [26]. Interestingly we found that AR occupancy causes a high level of somatic mutations at the DNA binding sites. The mutations observed at these binding sites was very different than the remainder of the genome.

## RESULTS

### AR binding sites have a markedly higher rate of mutations in PCa

To investigate the impact of TF binding on non-coding somatic mutations, we initially quantified the mutational density at binding sites using WGS of primary PCa (n=196) from the Pan Cancer Analysis of Whole Genome (PCAWG). TF binding sites were obtained from ChIP-seq of a single prostate cancer cell line (LNCaP), as very few studies have been done using clinical samples. DNA hypersensitive sites (DHS) were included as a negative control, as DHS were shown to have a lower rate of somatic mutations due to increased access of DNA repair machinery [27]. When we compared the mutational rate at TF binding sites to randomly shuffled regions in PCa, many TF binding sites including HOXB13, EP300, SUZ12 were found to have a statistically higher rate of mutations (FDR=0; **Figure 1**). As expected, DHS had a much lower rate of mutations than either any other TF or random regions. Contrasting earlier work in both colorectal cancer [5] and melanoma [7], CTCF binding sites did not have an increased rate of mutations as compared to either random regions or regions nearby the TF binding site (**Supplementary Figure 1A**). However of all the TF’s characterized, AR binding sites were found to have the highest rate of somatic mutations. To confirm that was not an artifact of using binding site data from a cell line, we observed an even greater mutational rate at ARBS that were identified from ChIP-seq of clinical PCa samples (**Figure 1**). A similar trend was observed with indels at ARBS in PCa, though not as dramatic due to the low numbers of indels obtained by consensus mutation calling (**Supplementary Figure 1B**). The increase in ARBS mutations is not likely due to epigenetic modifications as ARBS had greater than twice the mutation rate of regions with H3K27Ac, H3K4me3, H3K4me1 or H3K36me3 marks.

**Figure 1:**
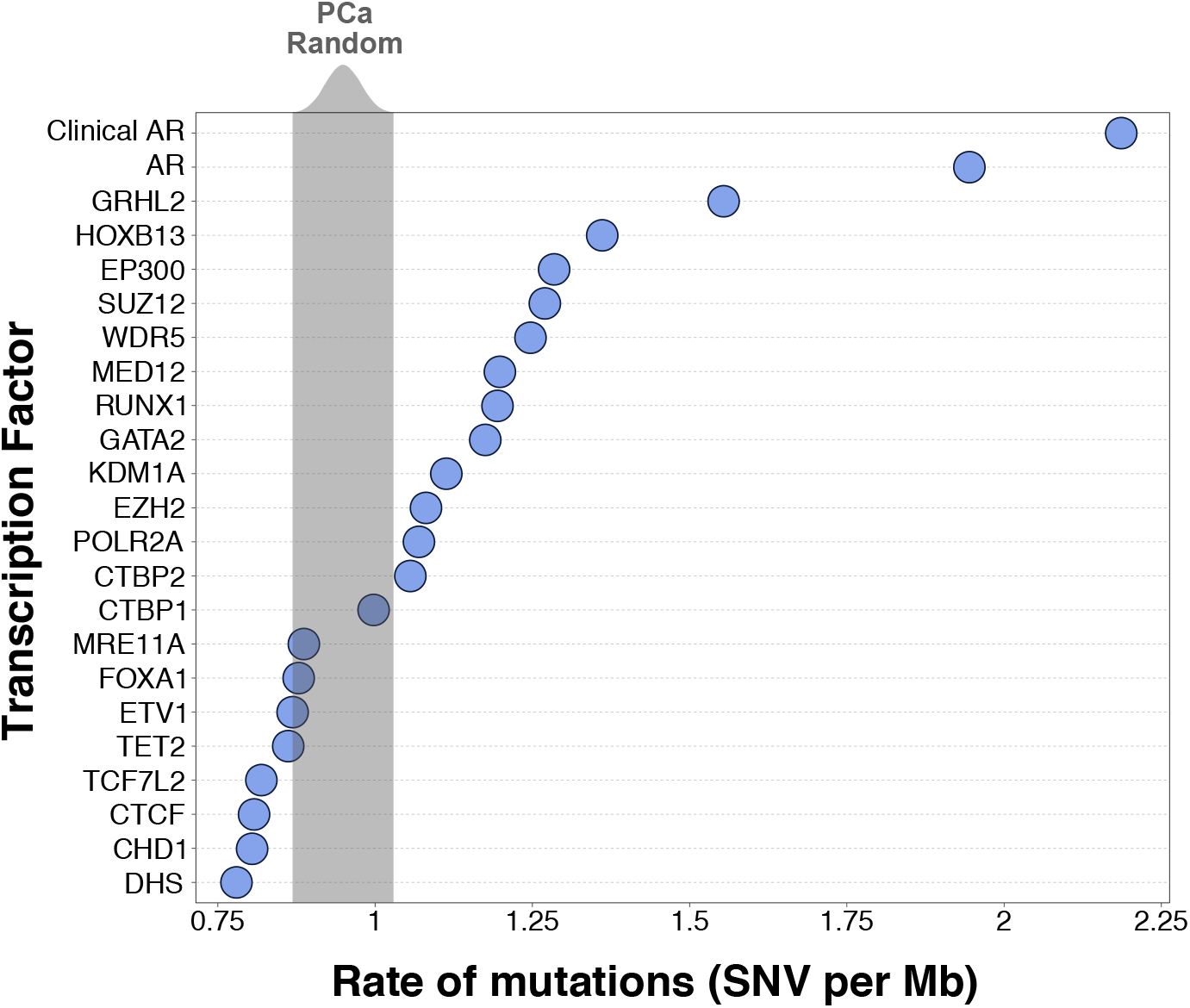
ARBS sites are the most heavily mutated TF binding sites. The rate of mutations (SNV per Mb) at individual TF binding sites (n=22) and DHS regions were compared to randomized chromosomal regions (1000 iterations, grey). All TF binding data was generated from a secondary cell line except “Clinical AR” that are high-confidence AR binding sites from patients PCa samples.

AR provides an ideal model to study TF-mediated mutations as this nuclear receptor is critical to the growth of nearly all PCa tumours, but is not active or required in other cancers. Thus, the ARBS chromosomal locations should not have increased mutations in other cancers if the observed results are due to AR binding rather than regional DNA instability. When we calculated the rate of mutations at ARBS from whole genome sequencing of over 20 different cancer types (n=2576) the rate of SNV mutations at ARBS was greater in PCa than either all other cancers (Wilcox t-test; *p<2x10^16^*) or any individual cancer (**Figure 2A**). Importantly, no other cancer other than PCa had a higher rate of SNVs at ARBS than random regions (**Supplementary Figure 2**). An increase in mutations at ARBS was clearly observed in PCa, but not other cancers, with a enrichment approximately ±375bp from the maximal AR peak (**Figure 2B**). This was not due to nucleotide composition, as those regions that have an ARE motif but no bound AR did not have an increase in SNVs or indels (**Figure 2C**). Providing further confidence that these mutations occur due to AR occupancy, we observed a clear correlation between SNV density and ChIP-seq peak height (**Figure 2C**). Overall these results demonstrate that AR binding correlates with an increase in somatic mutations.

**Figure 2:**
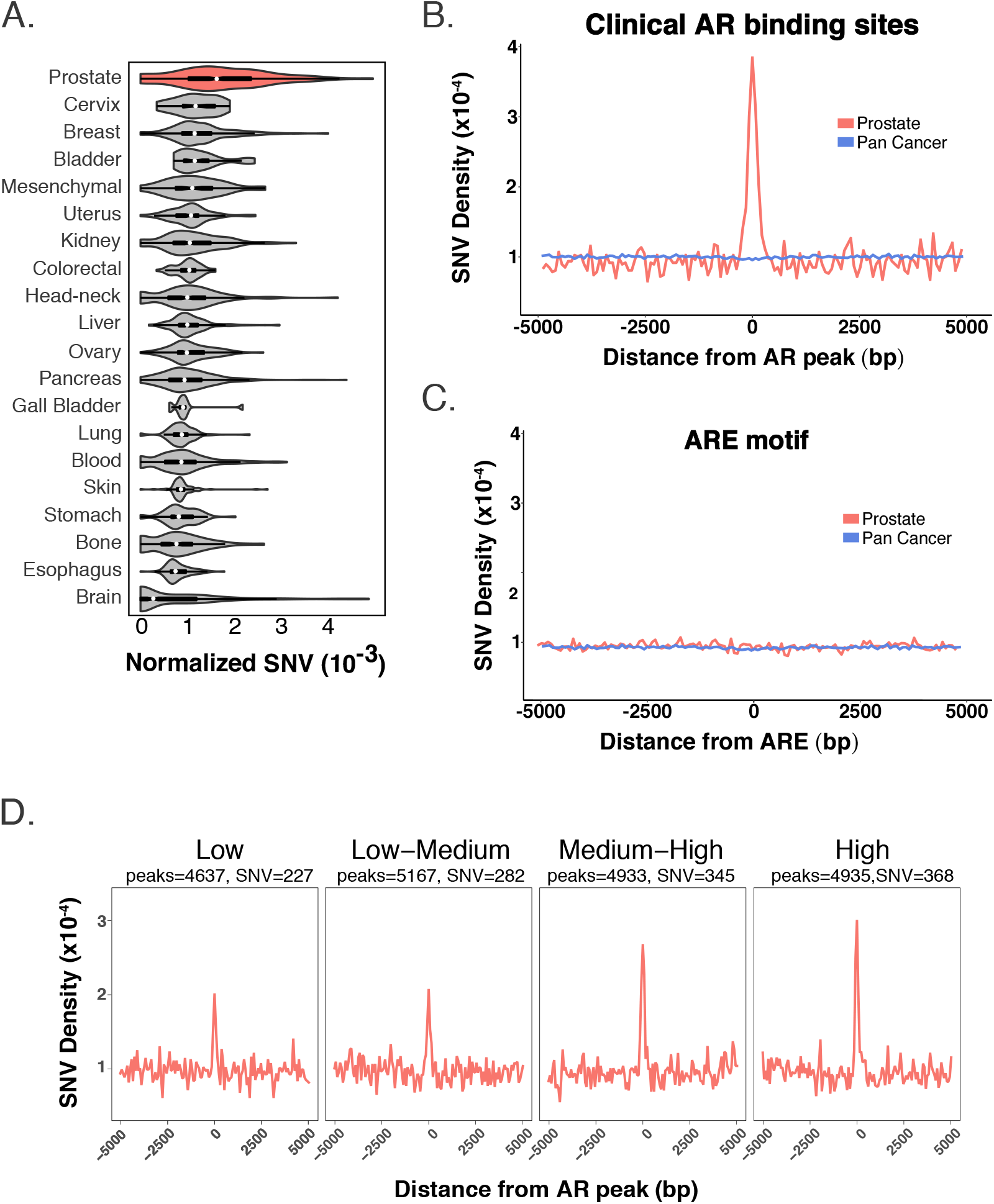
ARBS have an increased rate of mutations only in PCa. (A) PCa (red) has the highest normalized SNV rate at clinical ARBS of all cancer types. *(B)* The mutational density +/− 5kb at clinical ARBS was markedly increased in PCa (red) but not in all other cancers (blue). *(C)* A similar analysis was done at regions in the genome that had the canonical ARE motif but no AR binding. No increase in mutational rate was seen in either PCa (red) or other cancers (blue). (*D)* AR ChIP-seq peaks were divided into quartiles based on peak height (low/low-medium/medium-high/high). A clear correlation was observed between peak height and increased SNVs at ARBS.

To determine if a similar increase in mutations was observed with other lineage-specific TFs, we quantified the rate of SNV mutations at Estrogen Receptor binding sites (ERBS) in breast cancer (**Supplementary Figure 3**). Similar to what we observed at ARBS in PCa, breast cancer had the highest rate of mutations at ERBS. While not the goal of this work, it does suggest that TF binding site mutations are cell-of-origin specific.

We then looked to determine if the ARBS mutations occurred at those regions with specific epigenetic modifications or TF binding co-occupancy. This was based on previous literature which demonstrated that the cellular epigenetic state could dramatically alter the mutational rate [2]. However, despite extensive optimization no relationship could be observed between ARBS mutations and specific histone marks or TF co-occupancy (Supplementary Figure 4). As we do not have binding information for all possible histone marks or TF there may yet be an undiscovered correlation. However, our current data suggests that ARBS mutations do not correlate with specific epigenetic modification or proteins and are solely due to AR binding.

### AR-mediated SNV mutations induce purine transversions

To better understand the cause of these mutations, we then determined the mutational signature at ARBS. While these binding sites only represents a small portion of the total genome (~100Kb), the mutational signature of a large region should be roughly the same as the whole genome if there is a sufficient number of mutations. Supporting this, we found that random regions with a similar size or nucleotide composition to ARBS almost always had a near identical mutational signature to the PCa genome (**Supplementary Figure 5A**). Further, the number of SNVs observed at ARBS are well over the previously calculated minimum threshold to decipher a mutation signatures with >95% accuracy [28]. Interestingly, when we looked at the mutations at ARBS in PCa we found a dramatically different mutational profile than the remainder of the cancer genome (**Figure 3A**). Specifically, there was an increase in TpG->ApG and CpG->GpG purine transversions. These infrequent mutations occur at a much lower rate in the remainder of the PCa genome. Demonstrating that this was not due to the nucleotide composition, those regions of the chromosome that have an ARE motif but no AR binding did not have the same type of mutations (**Figure 3A**). When we shuffled chromosomal locations to match the nucleotide composition of ARBS and recalculated the mutational signature, no random regions were found to have mutational signatures comparably enriched for TpG ->ApG (**Supplementary Fig 5B**). We observed no difference in either the rate or type of mutation if the ARBS had a canonical ARE (Wilcoxon signed rank test *p=0.97;* Supplementary Fig 6). Finally, to test if the mutations were due to the specific chromosomal locations where AR binds we compared the mutational signatures at ARBS in all cancer types (**Figure 3B**). Only PCa was found to have a different mutation type at ARBS. All others cancers, which do not express or require AR, had the same signature at both the ARBS and whole genome. This demonstrates that the observed ARBS mutational signature was not caused by differences in nucleotide composition or chromosomal characteristics and is due to AR binding.

**Figure 3:**
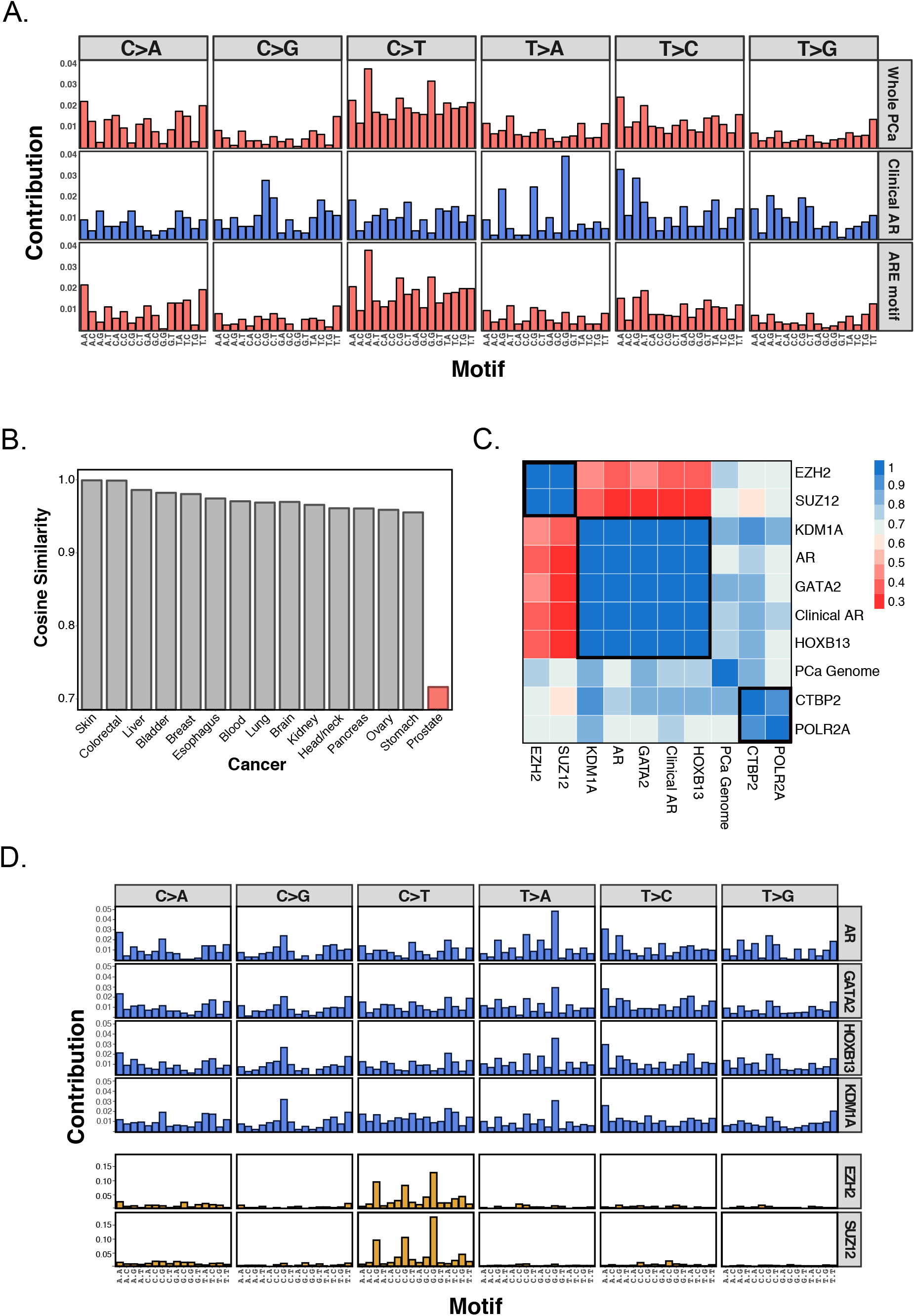
ARBS have a different mutation signature. (*A*) The type of mutations at clinical ARBS were compared to all those found in the whole PCa genome or those regions that contain an ARE motif but no AR. (*B*) The mutational signature at ARBS chromosomal regions was compared to the remainder of the genome in multiple cancer types. Only PCa was found to have an altered mutational signature at ARBS. (*C*) The mutational signature of each TFs binding site was compared and three clusters of mutation types were observed at SUZ12/EZH2 [group 1], KDM1A/AR/GATA2/HOXB13 [group 2] and CTBP2/POLR2A [group 3] (*D*). Detailed analysis of the group 1 and group 2 are shown.

Having observed an AR-specific mutational signature, we tested if other TF binding sites had similar types of mutations. If these were directly caused by AR binding, only ARBS would be expected to have this signature. We therefore analyzed all TFs that had both an increased rate of mutations (**Figure 1**) and a total number of mutations that was greater than the previously published theoretical threshold [28]. When the TF mutation signatures were compared, we found three distinct signature types (**Figure 3C**). First, KDM1A, HOXB13 and GATA2 were found to have a very similar mutational signature to AR (**Figure 3D**). This correlation was not due to an co-occupancy of the binding sites as a similar result was obtained even after removing regions that overlap with the AR (Supplementary Figure 7A+B). Further, it was not due to the nucleotide composition of these regions as those site with AR, GATA2 or HOXB13 motifs but no protein (motif alone) did not have either an increased rate of mutations or a change in the mutation type (**Supplementary Figure 7A+B**). These mutation types were only observed in PCa and were not seen in other cancer types (**Supplementary Figure 7C**). Second, members of the polycomb repressive complex 2 (PRC2) including SUZ12 and EZH2 had a very different mutational signatures than any other TF. As before, these mutations were not due to simple overlap of the binding sites (**Supplementary Figure 8A**). However, these specific mutations were not only seen in PCa. We also observed a similar mutational signature at SUZ12/EZH2 binding sites in several other cancer types (Supplementary Figure 8B). Finally, the remaining TFs including POLR2A and CTBP1 had a complicated mutational signature that was much closer to the whole genome than the other TFs. Importantly, the observed mutational signatures were not solely due to nucleotide composition as POLR2A, which has a similar GC content to SUZ12 and EZH2, had a very different mutational signature (**Supplementary Figure 9**).

To identify the potential etiological factor of the TF-mediated mutations we compared our results to previously published mutational signatures [1]. Demonstrating the utility of this method, there was a striking similarity between SUZ12/EZH2 binding sites and a previously published COSMIC mutational signature (*Signature 1*; **Figure 4A**). This well characterized signature has been reported in numerous cancer types and is caused by spontaneous deamination of 5-methylcytosine. Supporting this hypothesis almost all C->T mutations in SUZ12/EZH2 binding sites were found to occur at CpG sites (**Supplementary Figure 8C**). Further, when we looked at genome-wide bisulfite sequencing, SUZ12/EZH2 had some of the highest levels of DNA methylation of all the TF binding sites (Supplementary Figure 8D). Having shown the effectiveness of this approach, we then investigated the mutation type at ARBS and KDM1A/HOXB13/GATA2 binding sites. However, the ARBS mutations were very different than COSMIC mutational signatures and no published signature, excluding that caused by aristolochic acid, had a high frequency of TpG-> ApG mutations (**Figure 3A**). The aristolochic acid mutation signature (*Signature 22*) was excluded as it had many additional T->A mutations not observed at ARBS. Interestingly, the uncommon TpG-> ApG purine transversions have been previously shown to be caused by faulty repair of abasic sites. This is best demonstrated with the carcinogen dimethylbenzantracene (DMBA), which form a chemical adduct via one-electron oxidation that depurinates deoxyadenosine nucleotides [29–32]. This dramatic increase in abasic sites overwhelms the base excision repair machinery (BER), causing numerous TpG->ApG mutations due to the so-called “A-rule” whereby adenine substitutions are most likely to occur at unrepaired abasic sites [33]. It is not likely the prostate will be exposed to such polycyclic aromatic hydrocarbons, yet the same faulty repair could cause the ARBS mutational signature. Specifically, the presence of AR or other bound TFs could prevent the repair of spontaneously depurinated abasic sites. Such endogenous DNA lesions are extremely common and if not successfully repaired would cause the observed purine transversion. In support of this model, the ARBS mutational signature correlates well to that observed in DMBA treated animals (**Figure 4B**) [31].

**Figure 4:**
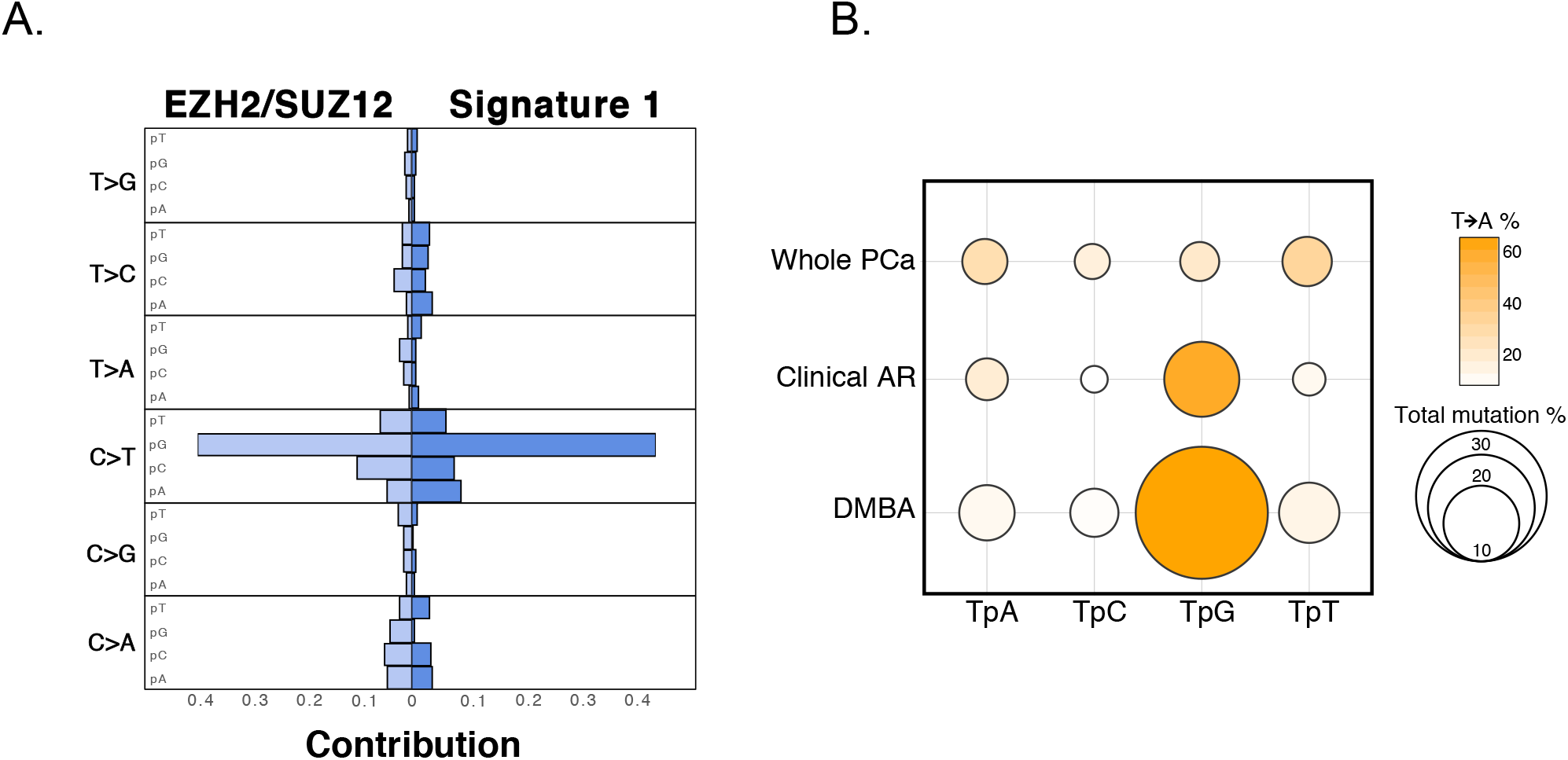
Mutation at PRC2 binding sites and ARBS are similar to previously published work. (*A*) EZH12 and SUZ12 mutations were concatenated into a PRC2 signature and compared to COSMIC signature 1. (*B*) The frequency and nucleotide composition of T->A transversions were compared between the whole PCa, clinical ARBS and DMBA treated mice. The majority of the the T->A mutation in both ARBS and DMBA treated mice were found to occur at TpG dinucleotides.

## DISCUSSION

Cancer is a disease of genetic errors. Through it’s mutations, we can begin to understand the molecular underpinnings of the malignant state. These mutations are not evenly distributed through the genome and are affected by numerous variables. There is emerging evidence that the rate of somatic mutations is higher at TF binding sites [5–7]. This has been demonstrated in two cancer types that have very high rates of mutations (~100 SNV/MB), as only a small number of patients is needed to provide sufficient statistical power. To determine if this phenomenon occurred in other cancers that have a lower rate of mutations, such as PCa, we used the recently released WGS from the PCAWG project. By working with an extremely large dataset of primary PCa (n=196) we could analyse how TF binding affected somatic mutations in this disease. We found that of all the TFs, ARBS had the highest rate of mutations with a clear correlation between mutation rate and AR occupancy. Importantly, an increase in ARBS mutations was not seen in those cancers that do not express or require AR. Some ARBS were highly mutated, with 2-3% of the patients having a mutation at these site. The high frequency of mutations at specific ARBS’s suggest that these mutations may modify the regulatory landscape of the cancer and potentially provide an evolutionary advantage. Interestingly, the type of mutations observed at ARBS was very different than those in the remainder of the PCa genome. Specifically, we saw a high frequency of TpG->ApG and CpG->GpG purine transversions at both ARBS and the binding sites of HOXB13, GATA2 and KDM1A. This is not due to overlap between the TFs, as the same type of mutations were observed at those binding sites that do not have AR cooccupancy. Previous research has demonstrated that AR activation can induce DSB by TOP2B, activation-induced cytidine deaminase or LINE-1 repeat-encoded ORF2 endonuclease [18,19]. While there was an increase in the rate of indels at ARBS, the type of mutations observed were not associated with DSB. In fact, compared to the remainder of the genome, ARBS had a decrease in the DSB mutational signature [1]. Our results suggest that DSBs that arise from AR-mediated transcription are efficiently repaired and do not cause a large number of SNVs. However, the impact of additional genotoxic stress, such as radiotherapy, on ARBS mutations requires further work. Of the AR pioneer factors, HOXB13 was found to have the highest rate of mutations. This supports recent work that demonstrated the AR cistrome of clinical PCa samples was reprogrammed from using FOXA1 to HOXB13 pioneer factors during tumourogenesis [9]. Supporting this, we observed a low frequency of mutations at the binding sites of FOXA1. This raises the interesting concept that TF mutational rate could be used as a surrogate for *in situ* activity. While speculative, the use of TF mutational rate may potentially help to identify potent pharmacological targets.

We observed a high frequency of mutations at SUZ12/EZH2 binding sites. Based on the type of mutations these were likely caused by 5-methylcytosine deamination. In support of this model, SUZ12/EZH2 had one of the highest rates of CpG methylation at TF binding sites. Similar mutations were observed at the same chromosomal locations in multiple cancer types. This suggests that these regions may be prone to this particular type of damage or, more likely, that PRC2 is important in these cancers. Regardless, the exclusivity of these mutation types at PRC2 binding sites raises many questions.

Mutations occur when repair of damaged DNA fails. Therefore, there are two potential mechanisms that could cause the observed increase in ARBS mutations. First, the AR itself may induce DNA damage when it binds to chromatin or induces gene transcription. Second, the bound protein may prevent access of repair machinery to the DNA lesion. While we cannot eliminate the first model, the increase in SNVs and remarkably similarity in mutational signature at several other TF binding sites including KDM1A, GATA2 and HOXB13, suggests that the AR itself does not induce DNA damage. Each of these TFs bind to a unique DNA sequences with different protein domains that function through disparate mechanisms. Such contrasting TFs are unlikely to induce similar damage. It is more probable that the increased rate of mutations is due to a blockade of DNA damage repair machinery. Supporting this, recent studies by Sabarinathan *et al*. demonstrated that in melanoma, TF binding impaired access of NER machinery [7]. By preventing the repair of UV-damaged DNA this led to a higher rate of mutations at TF binding sites. Our results in PCa support a similar, though expanded model. Specifically, we propose that TF binding prevents the repair of DNA by blocking not just NER but also BER. Several studies have demonstrated that TpG->ApG mutations, which were observed at ARBS, arise from the failed repair of abasic sites by BER. Abasic sites frequently occur due to spontaneous depurination at an estimated rate of 10,000 events per day per cell [34]. These endogenously damaged sites are typically repaired quickly and efficiently by BER machinery. While abasic sites can also be repaired by NER, this is much less common [35]. In our proposed model, TF binding prevents access of BER machinery to the damaged abasic sites. In support of this hypothesis, mice that have BER loss-of-function mutations accumulate endogenous DNA damage with increased rates ofs T->A and C->G purine transversions similar to that observed at ARBS [36]. Interestingly, the deamination-associated mutations observed at SUZ12/EZH2 binding sites are also caused by a failure of BER [37]. While NER has been shown to be impaired by TF binding to DNA, our results suggest that TF-mediated blockage may be a broader phenomenon that can impact other repair mechanisms.

Overall, this work demonstrates that somatic mutation distribution is influenced by lineage specific TFs. This complements previous studies and demonstrates that the cancer cell-of-origin influences mutation patterns. Given the critical role of AR in PCa and high frequency of mutations at ARBS, these mutations may affect cancer growth and development. However, further work is needed to understand how these mutations affect AR binding and gene transcription.

## METHODS

### Mutation Information of ICGC Patients

Whole genome sequencing data was obtained from Pan Cancer Analysis of Whole Genomes (PCAWG) release on August 24, 2016 [26]. For PCa only those patients with primary cancer (n=196) were included in the study due to the limited number of patients with metastatic or late-state prostate cancer. SNV and indels were previously called with three different mutation-calling algorithms (Sanger: indel=Pindel, SNV=Caveman; DKFZ: indel+SNV=Platypus; Broad: indel=Snowman, SNV=Mutect). Only those mutations which had been called by two or more callers and not found in dbSNP(v147) were used in this work.

### Transcription factor binding sites

ChIP-seq data was obtained for the following published work: FOXA1(GSM1410788), CHD1(GSM1573653), CTBP1(GSM1410762), CTBP2(GSM1410763), CTCF(GSM1006887), ETV1(GSM1145322), EZH2(GSM969570), GATA2(GSM941194), HOXB13(GSM1716764), MRE11A(GSM1543776), POLR2A(GSM1415124), RUNX1(GSM1527840), SUZ12(GSM969572), TCF7L2(GSM1249449), TET2(GSM1613322), TOP1(GSM1543792), WDR5(GSM1333369), KDM1A(GSM1279769), EP300(GSM686943), MED12(GSM686945), AR(GSE83860), H3K9ME3(GSM353610), H3K4ME1(GSM1410780), H3K4ME3(ENCODE: ENCFF401MDR), H3K27AC(GSM1249448), H3K36ME3(GSM875814). Clinical ARBS were identified from AR ChIP-seq of 13 tumour and 7 normal human tissue samples (GSE70079). Overlapping peaks were identified by HOMER’s (v4.7) *mergePeaks* function (−d parameter 200) [38]. All binding sites that overlapped with UCSD blacklisted regions were removed.

Motif driven peaks were predicted by PWMtools with given positional weight matrixes obtained from JASPAR DB.

### Determination of intersecting regions

Bedtools (version 2.26.0) and bedops (version 2.4.26) were used to intersect, manipulate and filter specific regions in bed and vcf files [39]. To extend binding regions bedtools *slop* function was used. For intersection and filtration, we used bedtools *intersect* and bedops *bedmap* function.

### Comparing specific region mutation frequency with background

Bedtools *shuffle* function was used to generate randomized regions across the genome. Each bed file was randomized 1000 times to generate a null distribution. All gapped regions (UCSC gapped regions) were removed. To generate random bed files with similar base composition (ATCG) of each random region we extensively randomized the AR binding data and then calculated base composition. We then z normalized each nucleotide type columns identify those random bed files similar to ARBS 250 bed file (as null value). The peak files which have the base composition that are in the ± 2 standard deviation (sd) range were selected.

### Mutation Signature Analysis

Mutation signature analysis was done using the bioconductor package SomaticSignature (version 2.12.1) with R version 3.4.0 [40]. Mutation signature were obtained from *plotMutationSpectrum()* function with default parameters. Those TFs with less than 480 mutations across all patients were not included in our analysis. This value was used as it was demonstrated to have a deciphering accuracy of >0.95 for two mutation signature [28].

As previously published, the *cosine()* function from the ‘lsa’ package was used to calculate the similarity between signatures obtained from SomaticSignature *motifMatrix()* function [28]

### Mutation aggregation analysis on TF binding regions

For each of the binding regions, overlapping mutations were mapped and mutation distances to the center of the TF binding region were calculated. For a given TF, each of the binding regions were overlapped based on their center. Mutation densities of 100 bp windows were calculated with smooth kernel density method. Calculation and visualization was conducted with ggplot2 R package.

### Methylation Analysis

CpG positions were identified from a published custom made perl script (https://www.biostars.Org/p/68352/#256983). DNA methylation was quantified from whole genome bisulfide sequencing from LNCaP cells (GSE86832). Methylation data points with coverage less than 10 were excluded from our analysis. Those locations with a DNA methylation less than 0.52 (median of LNCaP) were classified as unmethylated. Intersecting CpG of each peak was combined as a vector. Then all of the methylated and unmethylated sites were summed up to obtain single value of overall methylated rate of a TF. For given TFs, the intersection between TF and whole genome CpG was obtained.

### Heatmap

‘pheatmap’ package (version 1.0.8) was used for drawing heatmap from CRAN package repository *pheatmap* function was used in default settings to produce heat maps based on pairwise cosine similarity values of mutation signatures.

### Statistical Analysis

The distribution of mutation events limits the usage of parametric tests. For preventing biasing, we used R statistical language default *wilcox.test()* function is used for Wilcoxon rank sum test.

### Visualization

Data was visualized with *ggplot2* (version 2.2.1) and Venn diagrams were drawn in RShiny app, https://github.com/jolars/shiny-server.

## ACKNOWLEDGMENTS

We would like to acknowledge the ICGC/TCGA PCAWG consortium for providing access to this sequencing data. Further we would like to thank Eric LeBlanc for his comments on an early version of the manuscript and Firat Uyular for his computational suggestions. This work was supported by funding from Koç University School of Medicine and TÜBİTAK (114Z91).

## CONTRIBUTIONS

A.G. and O.K. analyzed data and provided computational support. M. G. analyzed data and provided statistical support, T.M and N.A.L. designed the experimental study, analyzed data and wrote the manuscript. All authors discussed the results and reviewed the manuscript.

## COMPETING INTRESTS

The authors declare no competing financial interests.

---

**Supplementary Figure 1:**

(*A*) SNV Density at CTCF binding sites in PCa. Unlike previous work in melanoma and colorectal cancer, CTCF binding sites have no increase of SNVs in in PCa. (*B*) Indel density in PCa and all other cancers (pan cancer). The rate of indels at ARBS was increased in PCa but not other cancers types.

**Supplementary Figure 2:**

The rate of SNVs at ARBS (blue arrow) was compared to a randomized region (grey) in all cancers with greater than 500 total SNVs. Only PCa (red) had a higher rate of mutations at ARBS than the null distribution.

**Supplementary Figure 3:**

The normalized rate of SNVs at ERBS was compared in multiple cancers. Breast cancer has found to have the highest rate of SNVs at these binding sites.

**Supplementary Figure 4:**

ARBS mutations were clustered with all other histone marks and TFs. No correlation was observed between the ARBS SNV and specific epigenetic marks or TF binding.

**Supplementary Figure 5:**

The mutational signature at ARBS is not due to nucleotide composition. (*A*) The mutational signature of PCa whole genome was compared against randomized regions (n=1000) that the same size as ARBS (~100k Kb). Randomized regions that had a matched nucleotide composition to the ARBS were subsetted (yellow). The mutational signature of the whole genome was extremely similar to the smaller randomized regions with a median cosine similarity of 0.953. (*B*) ARBS were found to have a higher frequency of all T->A or TpG->ApG transversion than randomized regions with a matched nucleotide composition.

**Supplementary Figure 6:**

The frequency and type of ARBS mutations were not impacted by the presence of a canonical Androgen Response Element (ARE) motif.

**Supplementary Figure 7:**

TF binding is required for an increase in SNV and altered mutation signature. (*A*) Mutation frequency was calculated at those sites that had a specific TF motif but no protein to those TF binding sites. Those regions which had the motif but no protein did not have a greater rate of mutations than randomize regiosn in the genome (grey)(*B*) Mutational signatures at motif containing and TF binding sites were calculated and analyzed using a cosine similarity measure. Motif signatures and ChIP-seq signatures appeared in two distinct clusters with the regions containing the motif being very similar to the whole PCa. (*C*) The cosine similarity of somatic mutations that occur at specific TF binding sites and the remainder of the cancer genome were calculated for all cancer types. Only PCa had a different mutational signature at these TF binding sites.

**Supplementary Figure 8:**

(*A*) EZH2 and SUZ12 ChIP-seq peaks shows a relatively low numbers of overlapping regions in LCNaP cells. (*B*) The mutation signature at SUZ12 and EZH2 binding sites is very different than the remainder of the genome in multiple cancer. (*C*) EZH2/SUZ12 mutation signatures were calculated at regions with CpG and non-CpG. C->T mutations were significantly enriched at CpG regions. (*D*) Overall methylation rate at TF binding sites was determined from whole genome bisulfite sequencing.

**Supplementary Figure 9:**

The GC and AT nucleotide composition was calculated at all TF binding sites, histone marks and regions that contain specific TF motifs.

